# Wetting of junctional condensates along the apical interface promotes tight junction belt formation

**DOI:** 10.1101/2022.12.16.520750

**Authors:** Karina Pombo-García, Cecilie Martin-Lemaitre, Alf Honigmann

**Affiliations:** Max Planck Institute of Molecular Cell Biology and Genetics, Dresden, Germany; Technische Universität Dresden, Biotechnologisches Zentrum, Center for Molecular and Cellular Bioengineering (CMCB), Dresden, Germany; Cluster of Excellence Physics of Life, TU Dresden, Dresden, Germany

## Abstract

Biomolecular condensates enable cell compartmentalization by acting as membrane-less organelles^1^. How cells control the interactions of condensates with other cellular structures such as membranes to drive morphological transitions remains poorly understood. Here, we studied formation of tight junctions, which initially assemble as condensates that over time elongate around the membrane cell perimeter to form a closed junctional barrier^2^. We discovered that the elongation of junctional condensates is driven by a physical wetting process around the apical membrane interface. Using temporal proximity proteomics in combination with live and super-resolution imaging, we found that wetting is mediated by the apical protein PATJ, which promotes adhesion of condensates to the apical membrane resulting in an interface formation and linear spreading into a closed belt. Using PATJ mutations we show that apical adhesion of junctional condensates is necessary and sufficient for stable tight junction belt formation. Our results demonstrate how cells exploit the collective biophysical properties of protein condensates and membrane interfaces to shape mesoscale structures.

## INTRODUCTION

Tight junctions are supramolecular adhesion complexes that control the para-cellular flux of solutes by forming diffusion barriers between cells ^3–5^. Junctional assembly is initiated by condensation of cytosolic scaffold ZO proteins at the cell-cell contact sites that over time elongate and fuse around the apical cell perimeters into a continuous belt which seals the tissue^2,6,7^. How the initially round junctional membrane condensates change composition and shape into a continuous tight junction belt is unclear. Interactions of biomolecular condensates with cellular structures such as membranes can, in principle, drive morphological changes via forces emerging at the interfaces of condensates^8^. Depending on the force balance of surface tensions and membrane adhesion strength, condensates can either give up their spherical shape by spreading on the surface of the membrane, akin to a water droplet wetting a hydrophilic surface, or membranes can bend around round protein condensates^9–11^. How cells tune the material properties of junctional condensates, such as in other cases ^12–14^, to control the forces created at the interface ^15,16^ and drive tight junction formation is not understood. Here we uncovered an example how cells use and modulate interfacial forces of membrane attached condensates to position and shape cell-cell adhesion complexes. Interestingly, these emergent morphological wetting transitions are directly related to the molecular composition of condensates and the membrane, which offers molecular control how cells can tune condensate shapes in space and time.

### Time-resolved proximity proteomics reveals the molecular composition of junctional condensates during tight junction belt formation

To understand how the tight junction belt is assembled and positioned, we combined APEX2 proximity proteomics labelling of the main junctional scaffold protein ZO1 with a calcium switch tissue formation assay in living epithelia cells^17,18^ (Figure 1A and Supplementary Figure 1A). Time resolved APEX2 has been instrumental in systematically dissecting the molecular composition of condensates due to its fast kinetics ^19,20^. Combining APEX2 protein profiling with a calcium switch assay allowed us to synchronize the initiation of junction assembly in the entire tissue by the addition of calcium to the culture medium and quantify the time evolution of the junctional proteome during the assembly process using proximity proteomics.

**Figure 1.**
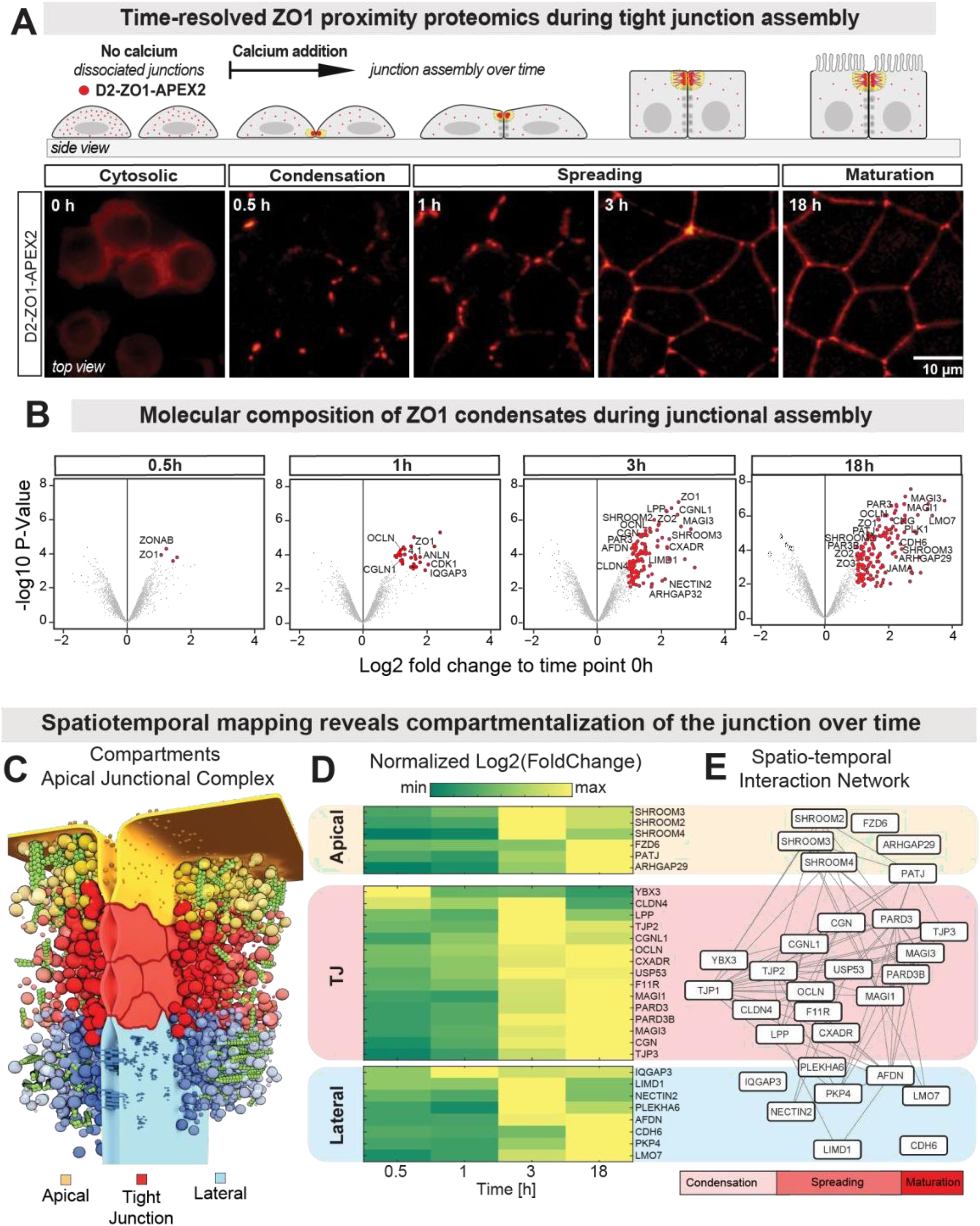
Time-resolved proximity proteomics reveals arrival times of proteins during tight junction belt formation. (A) Schematic representation of tight junction assembly using a calcium switch assay. Bottom live - imaging of MDCK-II cells labelled with Dendra2-ZO1-APEX2 after calcium switch, showing the morphological stages of tight junction belt formation. (B) Volcano plots displaying the log2 fold-change (FC) of biotinylated proteins around ZO1 as function junction assembly time against p-values calculated from the two biological repeats. All FCs were calculated against the initial time-point. Proteins with false discovery rate (FDR) < 0.05 (moderated t test) and FC ≥ 2 were considered significantly enriched and are depicted in red. Selected proteins are annotated. See Figure S1 for setup and controls of APEX2 assay. (C) Model of the apical junctional complex showing three membrane compartments, apical (yellow), tight junction (red) and lateral (blue). (D) Time resolved heat map of selected proteins of the apical, TJ and lateral compartments during junctional assembly. FC values are normalized. (E) Spatio-temporal protein interaction network relating the known position of proteins (apical, junctional, lateral) to the FC kinetics revealed by the APEX2-proteomics.

Fluorescence imaging of stably expressing Dendra2-ZO1-APEX2 in MDCKII kidney epithelia monolayers confirmed correct tight junction localization of the fusion construct (Figure 1A and Supplementary Figure 1A-C). Next, we confirmed the proximity labelling activity of the APEX2 enzyme by incubating the tissue with biotin-phenol for 30 min and subsequently induced the proximity labelling reaction by a short 1 min pulse of hydrogen peroxide (Supplementary Figure 1A). After quenching, fixation and staining with fluorescent streptavidin, two color imaging showed that the biotinylated proteins were highly enriched at the tight junction zone (Supplementary Figure 1C), demonstrating that Dendra2-APEX2-ZO1 proximity biotinylating provides high spatiotemporal contrast for mapping tight junction assembly. We further confirmed the APEX2 activity by blotting the biotinylated pull-down proteins (Supplementary Figure 1D).

Synchronized tight junction assembly was induced by sudden restoration of physiological calcium levels after overnight calcium depletion (Figure 1A). In line with previous studies, we found that formation of a closed ZO1 belt surrounding each cell took around 3h to 5h (Figure 1A, Video 1) ^2,21^. Based on the dynamics of ZO1 distribution on the membrane we subdivided the tight junction assembly process into 4 morphological stages. Between 0h – 0.5h ZO1 started to condense on the membrane of nascent cell-cell contacts. Between 1h – 3h ZO1 condensates spread and fused into a continuous belt. Finally, between 3h – 18h the junctional belt and cell shapes equilibrated into a stable confluent epithelial monolayer. To capture to molecular changes that go along with the morphological transitions, we performed ZO1 proteomics proximity-labelling at 0h, 0.5h, 1h, 3h and 18h after calcium switch, by activating APEX2 with a 1 min pulses of hydrogen peroxide, followed by isolation of the proteins via streptavidin pull down and digestion into tryptic peptides (Supplementary Figure 1E). To process all time-points in the same quantitative mass-spec analysis, we used a multiplex proteomic approach based on tandem mass tag (TMT). The TMT isobaric tagging approach enables robust quantitative proteomics by measuring all samples in one run. Hence, statistical analysis of relative protein enrichment at different time-points after calcium switch was possible across proteins detected in all time-points (Supplementary Figure 1F-H).

To analyze the changes of the ZO1 interactome as a function of tight junction assembly time, we calculated the fold change (FC) of protein abundance with respect to the time point zero, i.e. the calcium depleted state with dissociated junctions and cytoplasmic localization of ZO1. Proteins with FC > 2 were considered as potential hits and a false discovery rate threshold of 0.05 was used to filter out noisy data. In addition, we excluded false positive hits due to non-junctional interactions (ribosomes, nucleus, mitochondria, ER). The results of the analysis are shown as volcano plots in (Figure 1B). Proteins that significantly increased (hits) with respect to the dissociated state are highlighted in red and selected tight junction proteins are annotated. The protein proximity analysis revealed that the junctional condensates increase in molecular complexity during the assembly process (Figure 1B). We found that the majority of the proteins appeared after ZO1 membrane condensates formed around 0.5h, showing that initial composition of ZO1 condensates is strongly remodelled over time. The most significant increase in junctional components occurred after 1h, which corresponds to the time when the tight junction belt starts to close in live cell imaging (Figure 1A).

Taken together, our ZO1-APEX2 proteomics proximity profiling provides the first quantitative map of protein arrival kinetics during tight junction assembly. We use this dataset as a resource to understand the molecular underpinning of tight junction belt formation.

### Recruitment kinetics reveal polarization of junctional condensates during belt formation

The tight junction is known to be compartmentalized into different functional regions, which have been suggested to be important for assembly and positioning (Figure 1C). Interactions with adherens junctions in the lateral membrane have been implied in initiation of junction assembly at cell-cell contacts and providing mechanical robustness^22–24^. Interactions of the tight junction with apical membrane are thought to be important for positioning and signalling of the junctional belt 25–27.

To understand when the interactions with the lateral and apical compartments are established, we grouped our time resolved proximity proteomics data according their known compartment identity and analysed their recruitment kinetics (Figure 1C-E)^28^. This analysis revealed that the first adhesion receptors that are enriched around ZO1 are claudin-4 and later nectin-2, JAM-A and occludin, which are classified as core tight junction proteins ^29–31^. Earlier work on junction assembly suggested that the nucleation stage of tight junctions involves the recruitment of ZO proteins to adherens junctions via alpha-catenin^23,32^. Interestingly, our time-resolved ZO1 interactome does not support this view. Our data rather suggest that interactions with cadherin complexes (CDH6) are established at a later stage possibly via afadin (3-18h). In addition, we found that the RNA-binding protein YBX3 (ZONAB), which regulates cell proliferation, is sequestered to ZO1 condensates very early (0.5h) but is gradually excluded from ZO1 interactions as the junction matures, indicating a rather transient role in signaling^33,34^. The majority of junctional scaffold proteins such as MAGI3, PAR3 and CGN are recruited into junctional condensates between 1-3h^26,35^. Interestingly, components of the apical compartment such as members of the SHROOM family which are cytoskeletal adapters involved in cell shape regulations and the polarity protein PATJ are recruited during the junction spreading phase (~3h)^25,36^. At the final time point (18h) the majority of interaction partners around ZO1 show a stable enrichment with a few exceptions, which abundance decreased again at the last stage (SHROOMs, CLDN4, NECTIN2).

Taken together, our spatio-temporal recruitment analysis established the composition and compartmentalization of junctional condensates during the early, intermediate and late phase of tight junction assembly. We found that after the initial ZO1 condensation around a core set of tight junction proteins, the junctional condensates polarize, e.g. make contact with apical components, around the same time when junctional condensates elongate around the cell perimeter. which suggests a functional connection between these morphological and molecular processes.

### Apical proteins have slow recruitment kinetics to the junctional condensates

To directly visualize and verify individual protein arrival kinetics during tight junction assembly, we used quantitative live imaging of endogenously tagged proteins identified in the proteomics analysis (Figure 1D). We chose one candidate from the early (ZO2), intermediate (MAGI3) and late (PATJ) stage and used CRISPR/Cas9 to create fluorescent reporters with mScarlet (mS), respectively (Supplementary Figure 2A). Fluorescent tagging of candidate proteins was done in a MDCK cell line which expressed mNeoGreen (mN) tagged endogenous ZO1, to enable 2 colour imaging of junction assembly. Sequencing confirmed homozygous insertion of the tags (Supplementary Figure 3A) and imaging confirmed that mS tagging of the endogenous proteins resulted in proper co-localization with mN-ZO1 at the tight junction belt in confluent monolayers (Supplementary Figure 2A).

Next, we performed live-imaging of the 2-Color cell lines using the calcium switch assay to quantify protein arrival kinetics during tight junction assembly for 3 hours with 1 minute time resolution (Supplementary Figure 2B). To determine the arrival kinetics of the mScarlet-tagged tight junction proteins we segmented the mN-ZO1 signal and quantified the mScarlet intensity in the segmented ZO1 condensates and the cytoplasm over time (Supplementary Figure 3B). To directly correct for photo-bleaching artefacts, we calculate the ratio between the junctional and the cytoplasmic mScarlet signal for each time point (Supplementary Figure 3C). Assuming that bleaching is spatially homogenous the ratio is independent of bleaching and directly reports the enrichment of the protein in the condensed ZO1 phase (tight junction). The kinetics of the junction enrichment ratio showed that mS-ZO2 was rapidly enriched after ZO1 condensation (Supplementary Figure 2C). In comparison, mS-MAGI3 was recruited slower and reached saturation later. Finally, mS-PATJ recruitment was delayed even longer and showed a visible enrichment only around the time of junction spreading (Supplementary Figure 2B, C). In order to calculate the half times (t1/2) of client protein arrival we fitted the kinetic data to a HillSlope model (Supplementary Figure 3C) and determined the difference in arrival time between ZO1 and the client protein in living cells. The analysis confirmed the trend of the recruitment kinetics observed in the proximity proteomics dataset with early ZO2 (~5 min), intermediate MAGI3 (~22 min) and late PATJ (~35 min) (Figure 2C).

**Figure 2.**
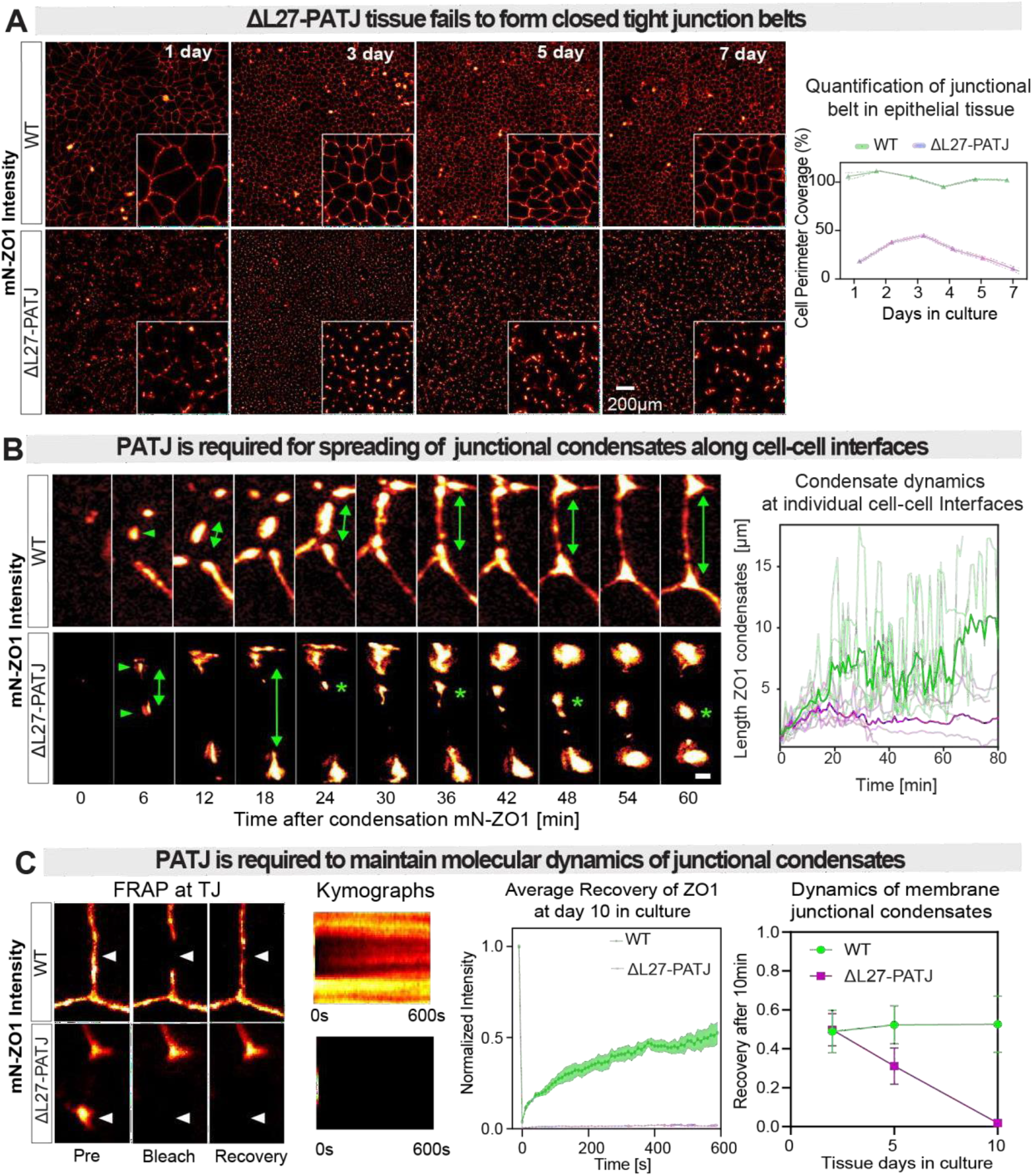
Spreading of junctional condensates along membrane requires PATJ. (A) Quantification of ZO1 belt formation in growing WT and ΔL27-PATJ MDCK-II tissue over 7 days after seeding. WT cells form a continuous and stable ZO1 belt from day 1. In ΔL27-PATJ tissue ZO1 remains fragmented even after 7 days of culture. Right panel shows a quantification of cell perimeter coverage (CellPose segmentation) coverage of mN-ZO1 condensates (local thresholding) in WT and ΔL27-PATJ tissue (see Figure S3B). In WT tissue ZO1 condensates cover 100% of the cell perimeters within the first day. In ΔL27-PATJ tissue cell perimeter coverage never exceeds more than 50% of the cellular perimeters and even decreases over time (n=3 experiments, mean ± SD). See Movie S1 and S2 for the dynamics of mN-ZO1 in the first hour after calcium switch in WT and ΔL27-PATJ tissue. (B) Analysis of ZO1 junctional condensate dynamics at single cell-cell interfaces after calcium switch. ZO1 condensates form at the cell interface (green arrows). Then elongate and fuse into a continuous belt in WT (double side arrow) (Movie S3). In ΔL27-PATJ cells condensates still form but fail to elongate and fuse MDCK cells (green stars) (Movie S4). Right panel shows quantification of the length of ZO1 condensates over time. Time 0 min is considered from the first ZO1-condenste on the membrane. In WT cells length of ZO1 condensates on average increases over time to a length of 10 μm (bold green). Single condensates show strong fluctuations of length due to transient fusion and fission events along the cell perimeter (light green). In ΔL27-PATJ cells the average length of ZO1 condensates remains below 5 um (bold pink) and the fluctuations due to elongation and fission are strongly reduced (light pink) (n=10). (C) Quantification of dynamics of mN-ZO1 in junctional condensates via FRAP. mN-ZO1 was bleached at the membrane, and recovery was measured at room temperature over time. Kymographs show that ZO1 recovered rapidly from the cytoplasm in WT. Reduced recovery was observed in ΔL27-PATJ cells indicating that dynamics of ZO1 in condensates are slower. Averaged recovery curves of 10 independent measurements (mean ± SD) are shown. Right Panel shows the mobile fraction after 10min recovery over the course of several days.

Taken together, live imaging confirmed the protein recruitment kinetics established by time resolved APEX2 proteomics to the junction for selected proteins from the early, intermediate and late assembly phase. Importantly, we confirmed that the polarity protein PATJ arrives to the junction around the time of junctional condensate elongation, suggesting the interaction of ZO1 condensates with the apical membrane is important for the morphological transition of single condensates into the continuous junctional belt.

### Spreading of junctional condensates along the apical membrane interface requires PATJ

Previous studies have shown that PATJ is multi-domain scaffold protein that binds to the apical polarity crumbs-complex via its N-terminal L27 ^37^. PATJ depletion via RNAi has been shown to cause a delay in tight junction formation ^25,38^. To investigative the functional role of PATJ for spreading of ZO1 condensates and tight junction belt formation, we deleted the N-terminal L27 domain via CRISPR/Cas9 gene editing in the background of the mN-ZO1 knock-in MDCK-II (Figure 2A). Sequencing of two clones confirmed deletion of the first exon including a frame shift leading to an early stop codon (Supplementary Figure 4A, left panel). Western-blots and qPCR against the N-terminal L27 domain of PATJ, encoded by the first exon, confirmed deletion of this domain (Supplementary Figure 4A, right panel). Immunostaining of the more C-terminal PDZ4 confirm that the remaining truncated protein was located in the cytosol and was not present the tight junction (Supplementary Figure 4D).

Imaging of mN-ZO1 distribution in confluent ΔL27-PATJ monolayers revealed a severe disruption of the tight junction belt compared to WT tissue (Figure 2A). In contrast to the continuous junctional belts in WT tissue (Video 1), ZO1 formed isolated puncta at cell-cell contacts in ΔL27-PATJ tissue (Video 2). In line with this observation, quantification of the average belt length per cell and the normalized cell perimeter coverage of ZO1 in WT and ΔL27-PATJ tissue over the course of 10 days, showed an overall strong reduction of ZO1 belt length and perimeter coverage in ΔL27-PATJ cells (Figure 2A, Supplementary Figure 4B-C). In WT tissue the ZO1 belt covered close to 100% of cellular perimeters within the first day after seeding. In contrast, ΔL27-PATJ initially made fragmented ZO1 belts, which over time further degraded to a cell perimeter coverage of only 10 %. In line with the loss of continuous ZO1 belts, there was a loss of barrier function confirmed by the trans-epithelial permeability measurements with fluorescent dextran (Supplementary Figure 4E).

To understand what step in the tight junction assembly process was disrupted by PATJ truncation, we studied the difference in the dynamics of ZO1 in ΔL27-PATJ during junction assembly (Figure 2B). As observed previously, in WT tissue ZO1 condensates initiated at cell-cell contacts sites and then spread along the cell-cell interface and fused into a continuous junctional belt (Figure 2B, upper panel, Video 3). Quantification of the elongation of junctional condensates at single cell-cell interfaces showed that in WT tissue elongation on average increased over time and saturated at 10 μm/cell after 60 minutes (Figure 2B, dark green), which coincided with the closing of the belt seen in the images. We observed large fluctuations of condensate length seen in the single junction traces (light green). These fluctuations were caused by ZO1 condensates fusing and rupturing multiple times until a stable belt was formed. Thus, in WT tissue belt formation is a very dynamic process in which elongation and fusion of ZO1 around cell perimeters is challenged by mechanical extension of the cell-cell interfaces during tissue formation. Comparing the elongation process in the WT to ΔL27-PATJ tissue, we found that initial condensation at nascent cell-cell adhesion sites was hardly affected by ΔL27-PATJ (Figure 2B, lower panel) (Video 4). However, the loss of PATJ from the membrane inhibited the spreading and fusion of junctional condensates along the cell-cell apical interface, i.e. junctional condensates remained more spherical during the formation of the tissue reducing the average condensate length to 3 μm/cell (light pink) (Figure 2B, magenta). In many cases we observed that junctional condensates after initial growth even shrunk in size or completely dissolved. In addition, the junctional fluctuation dynamics at individual cell-cell interfaces showed a strong reduction compare to WT tissue. Together, the reduced spreading dynamics of ZO1 condensates in ΔL27-PATJ cells prevented a closure of junctional belt around cellular perimeters (Supplementary Figure 2C).

To check whether the reduced spreading of junctional condensates could be linked to a change in its material properties, we performed fluorescence recovery after photo-bleaching (FRAP) (Figure 2C). Interestingly, we found that ZO1 condensates in ΔL27-PATJ cells recovered significantly slower compared to WT cells as the tissue reached a post-confluent state (Figure 2C). This indicates that the molecular dynamics of junctional condensates are strongly reduced, which is in line with a more solid-like state of the condensates.

Taken together, the quantification of ZO1 condensate dynamics in WT and ΔL27-PATJ tissue, revealed that PATJ is required for the spreading and coalescence into a continuous, dynamic and functional tight junction belt. These results suggest that belt formation around the cell perimeter could be directly facilitated by interactions of ZO1 condensates with the apical membrane. In this scenario PATJ would act as a linker that mediates adhesion of ZO1 condensates to the apical membrane. Adhesion of liquid droplets to surfaces is known to drive wetting transitions, which result in spreading on the surface ^8,39^. Therefore, we speculated whether PATJ mediated adhesion could drive the elongation of ZO1 condensates along the apical interface by a wetting-like process.

### PATJ recruitment to junctional membrane condensates drives the spreading process

To test the idea if PATJ directly mediates the spreading and fusion process of junctional condensates, we performed fast 2 colour imaging of endogenous mN-ZO1 and mS-PATJ in epithelia tissue quantifying the dynamics of both proteins with 1 minute resolution (Figure 3A) (Video 5-6). In line with our observations in Figure 2, we found that PATJ was recruited to ZO1 condensates at the time of spreading along the cell perimeter. To quantify this process, we measured how the extension of junctional condensates relates to the enrichment of PATJ in the condensates (Figure 3B). The analysis revealed a strong positive correlation (R = 0.7) between the extension of condensates and PATJ enrichment within ZO1 condensates, i.e. PATJ was not detectible in small, round condensates but became significantly enriched as the condensate elongated (Figure 3B). In addition, plotting the time evolution of the average eccentricity of junctional condenses together with the junctional enrichment of PATJ clearly showed that both properties follow the same kinetics, i.e. round ZO1 condensates deform into an eccentric elongated shape at the same time as PATJ concentration increases (Figure 3C). Taken together, the strong correlation between elongation of ZO1 condensates and PATJ enrichment support the idea that PATJ is directly involved in the condensate spreading. In addition, the similar kinetics of elongation and PATJ enrichment suggest that arrival of PATJ to ZO1 condensates initiates spreading.

**Figure 3.**
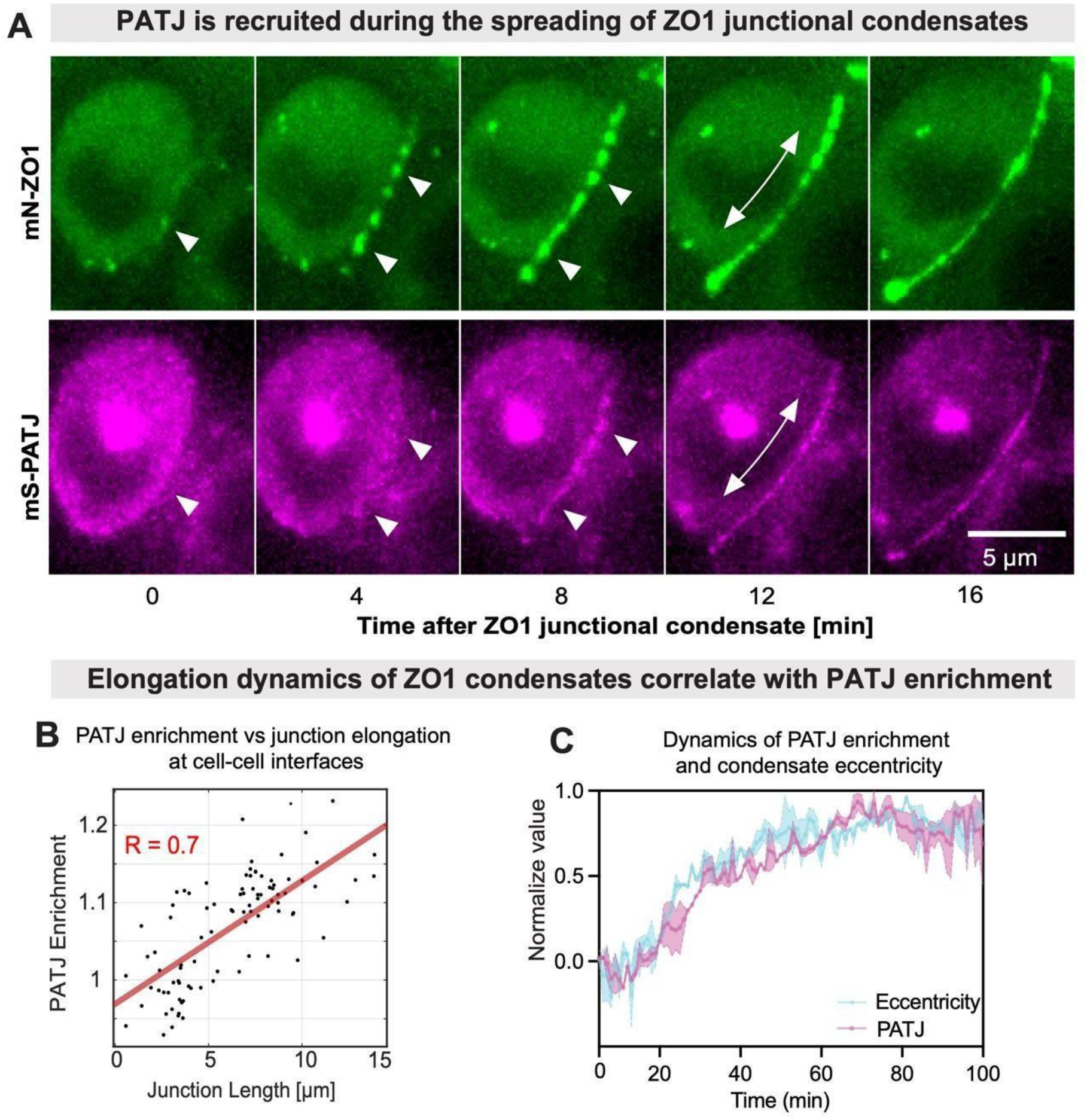
PATJ recruitment to junctional condensates correlates with spreading. (A) Live-imaging of double knock in of ZO1-mN and PATJ-mS cells showing the recruitment kinetics during the spreading of condensates. See also Movie S5 and S6. (B) Quantification of PATJ enrichment in ZO1 condensates vs elongation of condensates reveal a strong correlation between PATJ enrichment and condensate elongation (R=0.7). (C) Quantification of PATJ enrichment and condensate eccentricity changes over time show that both processes follow a similar kinetic. Error bars SEM of 5 independent measurements.

### PATJ forms the apical interface of the tight junction belt

To better understand how PATJ promotes the spreading of junctional condensates along cell-cell interfaces, we performed 2-colour STED microscopy to determine the ultra-structure of PATJ and ZO1 at the apical cell-cell interface. To this end we used our recently developed combination of 2D STED microscopy with 3D tissue culture, which enables imaging cell-cell interfaces in the high-resolution plane (XY) of the microscope (Figure 4A) ^40^. STED imaging of fully formed junctions revealed that ZO1 formed a condensed belt at the apical cell-cell interface (Figure 4B). Towards the lateral side the belt ZO1 formed a network like structure reminiscent of tight junction strands observed by freeze fracture electron microscopy^41^. Interestingly, PATJ was strongly enriched at the apical interface of the condensed ZO1 junctional network, forming clusters around the most apical strand of the ZO1 network, in line with previous observations ^28,6^. Quantification of the nearest neighbour distance between the most apical ZO1 strand and PATJ clusters (Figure 4C) revealed a mean distance of 40 nm (Figure 4D). Thus, PATJ is in close proximity to ZO1, but it is locally excluded from the core of the ZO1 condensate in mature tissue.

**Figure 4.**
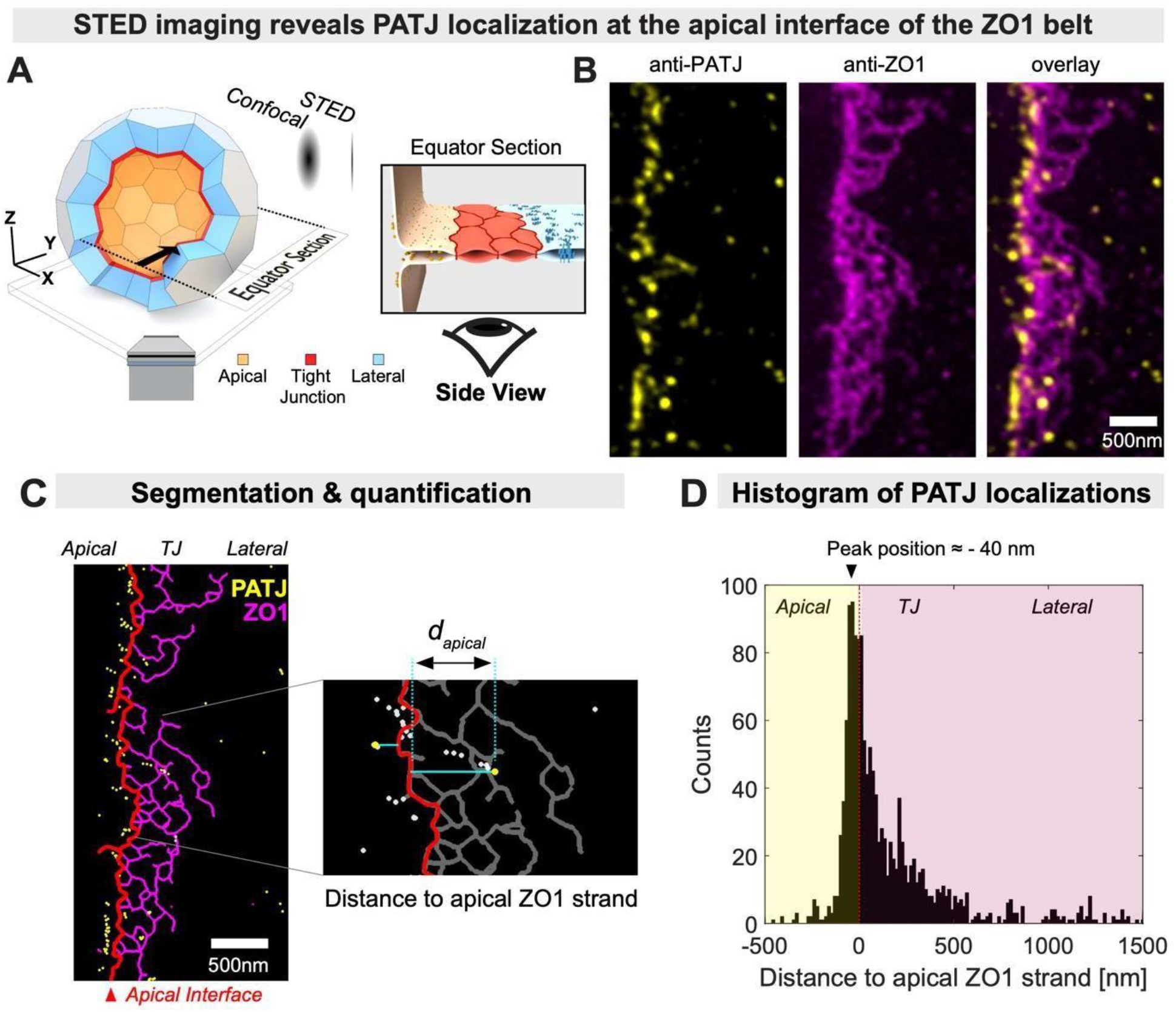
PATJ localizes to the apical interface of ZO1 condensates. (A) Combining 3D tissue culture of MDCK-II cysts with STED microscopy enables super-resolution imaging of cell-cell interfaces. At the equator cell-cell interfaces are oriented parallel to the high-resolution axis (XY) of the microscope. (B) 2-colour STED imaging of PATJ (yellow) and ZO1 (magenta) in MDCK-II cysts reveals that ZO1 forms a network structure reminiscent of tight junction strands. PATJ is enriched as clusters at the apical interface of the ZO1 network. (C) Segmentation of the ZO1 network and PATJ clusters enable quantification of the distance of PATJ with respect to the apical interface (apical ZO1 belt, shown in red). (D) Histogram of the distances of PATJ to apical interface shows a strong enrichment close to the apical interface.

Taken together, the super-resolution analysis revealed that PATJ forms the interface between the apical membrane and the ZO1 network, which supports the idea that PATJ directly mediates adhesion of ZO1 condensates to the apical membrane interface. In addition, the distance analysis between PATJ and ZO1 showed that PATJ is in very close proximity to ZO1 but remains locally excluded from the core of the condensate. This observation suggests two types of interactions underlying complex formation between ZO1 and PATJ. One adhesive interaction that mediates co-localization and one repulsive interaction that drives segregation. Based on previous work, we speculated that the adhesive interaction could be facilitated by short range binding of membrane bound PATJ to ZO proteins^25,28,42–44^. Repulsion could be the consequence of PATJ being part of the apical polarity mechanism via binding to the Crumbs complex, which would preventing mixing with the lateral membrane ^45,46^. The combination of long-range segregation between lateral and apical domains with short range binding interactions could provide a mechanism that leads to accumulation and spreading of junctional condensates along the apical-lateral interface.

### Adhesion of condensates to the apical interface is sufficient to rescue the belt formation

Our data so far indicate that ZO1 and PATJ form an interface that promotes the formation of a functional tight junction belt. To test the hypothesis whether a key function of PATJ is to promote condensate spreading by providing adhesion of ZO1 to the apical interface, we performed a structure-function analysis of the protein-protein interaction domains of PATJ (Figure 5A). Previous work on PATJ identified domains which mediate the interaction with apical Crumbs complex and putatively the ZO scaffold ^25,38^. Based on this data, we investigated the rescue of junctional belt formation and functional permeability of a series of truncations in the ΔL27-PATJ cell line (Figure 5).

**Figure 5.**
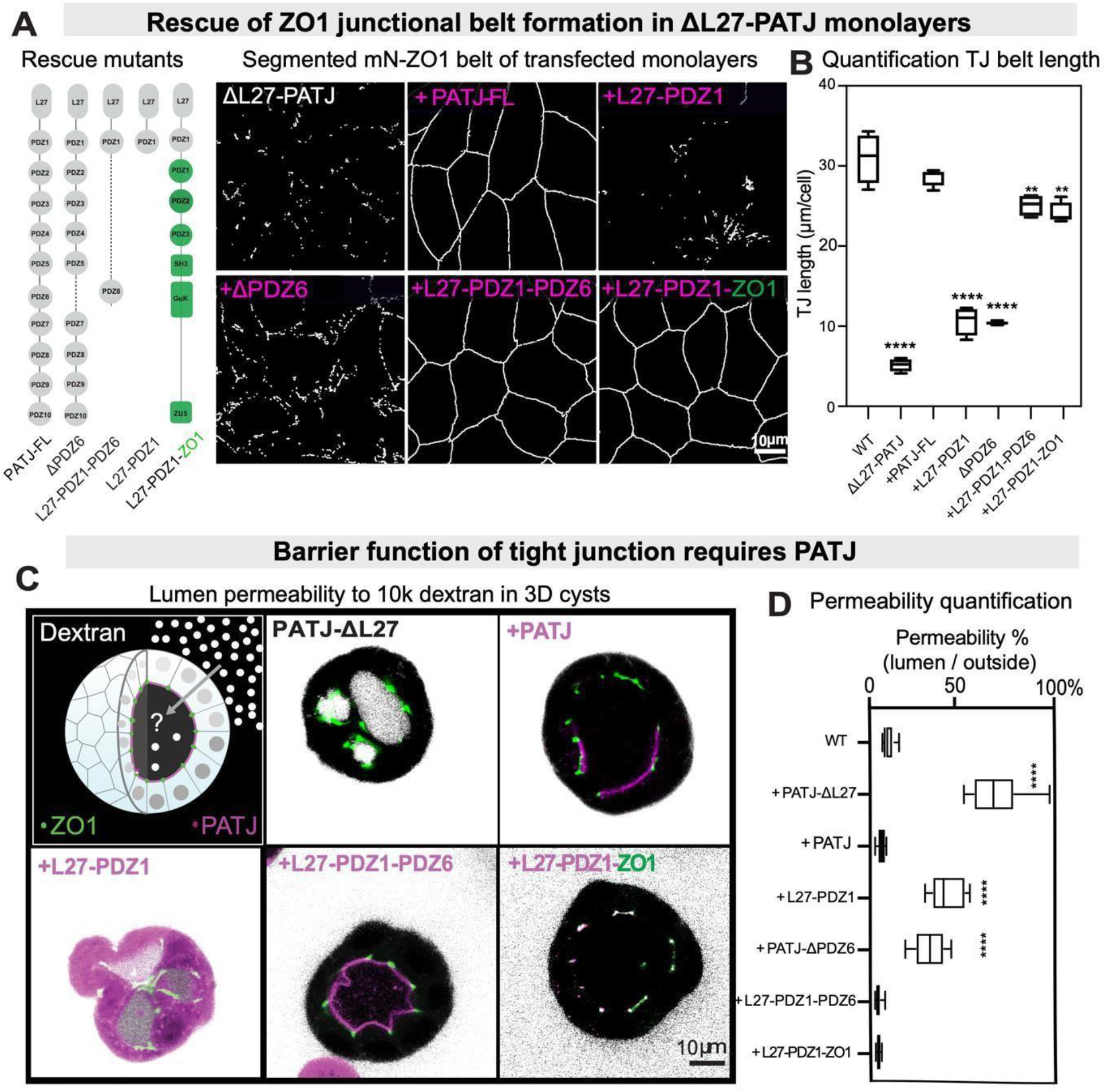
Adhesion of junctional condensates to the apical interface is required for belt formation and barrier function of the tight junction. (A) Scheme of PATJ mutants introduced into ΔL27-PATJ cells to rescue tight junction belt formation (B) and trans-epithelial permeability (C,D). Images show segmented maximum projections of confocal stacks of mN-ZO1 of MDCK-II ΔL27-PATJ transfect with the rescue constructs. The junctional belt was fragmented in ΔL27-PATJ and ΔPDZ6-PATJ and L27-PDZ1-PATJ monolayers, but restored in PATJ-FL and L27-PDZ1-PDZ6 and L27-PDZ1-ZO1 mutants. (B) Quantification shows the tight junction length/cell for the different rescue constructs (n>50 cells +/-SEM, asterisks indicate significance test with WT tissue using t-test ****P-value <0.0001. (C) Trans-epithelial permeability of ΔL27-PATJ MDCK-II cysts transfected with PATJ rescue constructs. 3D cyst grown in matrigel were incubated with dextran (10k-alexa647). Shown are confocal middle planes through 3D cyst made from WT (ZO1-mN KI) and ΔL27-PATJ cells expressing mCh-PATJ mutants. (D) Quantification of trans-epithelial permeability. Boxplot shows min and max □ SD, asterisks indicate significant comparison with the WT ZO1mN using a one-ANOVA test of n=10, ****P-value <0.0001.

To quantify the ability of PATJ truncations to rescue tight junction formation we determined the average ZO1 junctional belt length per cell (Figure 5B) and the barrier function of the belt by measuring the trans-epithelial permeability to 10k fluorescent dextran in MDCK cysts (Figure 5D) in ΔL27-PATJ MDCK tissue rescued with different PATJ constructs. As shown above, ZO1 belt length as well as the permeability barrier were severely compromised in the ΔL27-PATJ tissue compared to WT tissue. Expression of full length PATJ in the ΔL27-PATJ background resulted in full rescue of the belt length and the permeability barrier to WT levels (Figure 5B-F). Bringing back only the N-terminal L27-PDZ1 domains, which promote the interaction with the apical crumbs complex via Pals1 did not rescue belt formation nor the permeability barrier (Figure 5B-D). Similarly, PATJ lacking only the PDZ6 domain, which had been shown to be important for interactions with tight junction proteins was not able to completely rescue tight junction formation and barrier function ^38,47,48^. However, the expression of the Nterminal apical binding domain together with the tight junction interaction domain L27-PDZ1PDZ6 resulted in rescue of ZO1 belt formation and restoration of the permeability barrier (Figure 5C-D). This data suggests that PATJ requires both apical as well as tight junction binding capability to bring ZO1 to the apical interface and promote formation of continuous tight junction belts.

Since it is not clear whether PATJ interacts directly or indirectly with the ZO1 scaffold, we made a chimeric construct with the N-terminal apical binding module of PATJ (L27-PDZ1) fused to full length ZO1. Thus, we genetically provided ZO1 with the ability to interact with the apical membrane. Expression of this chimeric construct in the ΔL27-PATJ background resulted in rescue of the permeability barrier to WT levels and ZO1 belt length close to WT level (Figure 5BF). Interestingly, the localization of the chimera was similar to endogenous PATJ, apical and subapical (Figure 5C), not like the endogenous ZO1 localizing exclusively subapical. The PATJ truncations L27-PDZ1 and □L27-PDZ1, which did not rescue belt formation, were both distributed mainly in the cytosol. In contrast, all constructs, which did rescue belt formation, were in apical juxtaposition to the ZO1 belt (Figure 5C).

We conclude that the PATJ rescue experiments support the hypothesis that ZO1 junctional condensates interact with the apical membrane via PATJ. The chimeric ZO1-PATJ construct shows that direct interactions between PATJ and ZO1 are sufficient for spreading of ZO1 condensates along the apical interface and formation of a functional tight junction belt.

## DISCUSSION

Our study provides direct evidence that tight junction belt formation is driven by active wetting of junctional condensates around the apical membrane interface. Wetting in physics describes the spreading of a liquid droplet into a thin film on a surface^39^. Similar to wetting of liquid droplets on a surface, we found that ZO1 junctional condensates spread around the apical-lateral membrane interface in a process mediated by PATJ. However, different to wetting of 3D liquid droplets onto a 2D surface, ZO1 condensates are 2D structures that form on the lateral membrane domain and then spread along the 1D interface of the apical-lateral membrane. Despite these geometrical differences to conventional wetting, we expect the same physical principles to be at play. Wetting transitions of liquids on surfaces are driven by adhesive interactions between the droplet and the surface that overcome the surface tension of the droplet^49^. Along the same lines, wetting of ZO1 junctional condensates along the apical interface requires adhesive forces to the apical membrane that overcome the surface or line tension of the condensates.

Using our time resolved proximity proteomics data, we revealed how the molecular composition of junctional condensates evolves during the junction assembly process (Figure 1). We found that apical proteins come in molecular proximity to ZO1 during the time when junctional condensates elongate and spread around the apical cell perimeter (Supplementary Figure 2). Further functional experiments revealed that apical polarity protein PATJ promotes the reshaping of isolated membrane condensates into functional belt (Figure 2). In earlier work PATJ was found as a scaffold protein of the apical polarity complex that is enriched at tight junctions. However, its exact function in junction assembly remained unclear^25,44,50^. In addition to earlier work, we found that deletion of the N-terminal domain of PATJ (ΔL27-PATJ), which mediates the binding to the apical membrane, inhibits spreading of ZO1 condensates and results in leaky tight junctions^37^. This suggests that PATJ mediated interactions of ZO1 condensates to the apical membrane are necessary for spreading of condensates and formation of a functional tight junction belt. In line with this hypothesis, our endogenous tagged two-colour quantitative live cell imaging showed that PATJ is recruited to ZO1 condensates when spreading is initiated and elongation of condensate on the membrane strongly correlates with PATJ enrichment (Figure 3). Our super-resolution image analysis showed that PATJ is strongly enriched at the interface of ZO1 condensates and the apical membrane (Figure 4). Interface localization provides strong evidence that PATJ indeed directly mediates adhesion between ZO1 condensates and the apical membrane. Finally, we show that condensate spreading and tight junction belt formation can be rescued in ΔL27-PATJ tissue by a chimeric ZO1-L27 construct which directly provides ZO1 with ability to adhere to the apical membrane (Figure 5). Together, these experiments suggest that (i) PATJ mediates adhesion between ZO1 condensates and the apical membrane interface and (ii) apical adhesion is required for condensate spreading along the membrane and tight junction belt formation.

Based on our results we propose a model that explains formation of the tight junction belt as a consequence of wetting of junctional condensates around the apical cell perimeter (Figure 6). First, tight junction formation is initiated by ZO1 condensation at cell-cell contacts. This leads to formation of isolated junctional foci in the lateral membrane domain which sequester proteins required for assembly of tight junctions. Subsequently, junctional condensates interact with PATJ, which is anchored in the apical membrane via the Crumbs complex. Adhesion to the apical membrane induces a wetting transition of junctional condensates which drives spreading along the apical interface and fusion into a continuous functional belt.

**Figure 6.**
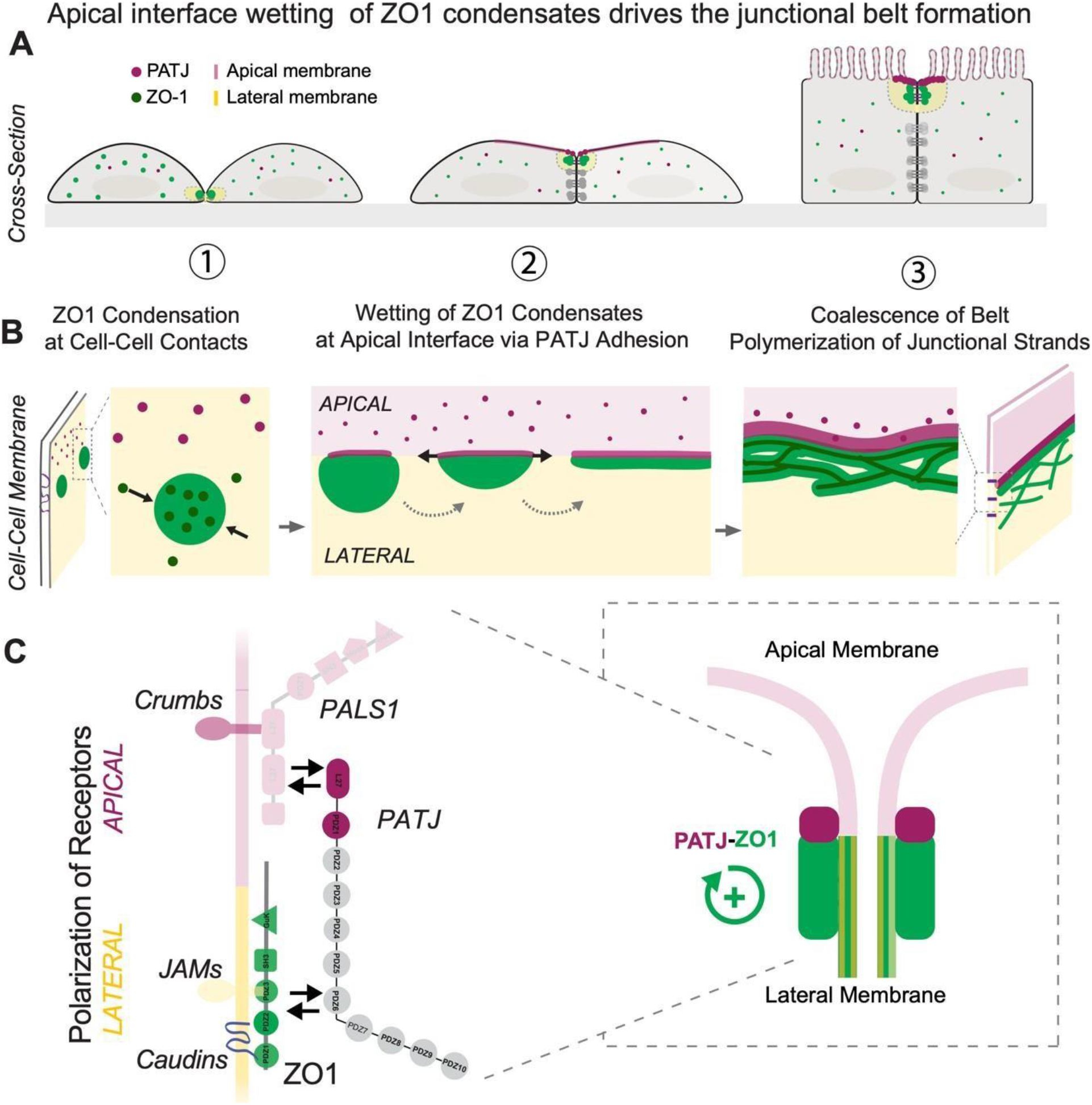
Model of the junctional belt formation by wetting of junctional condensates along the apical membrane interface. (A) Cross-section view of tight junction formation on the cellular level. (1) ZO1 (green) condensates form at cell-cell contact sites. (2) Cells polarize and ZO1 condensates adhere to the apical membrane via interactions with PATJ (magenta). ZO1 condensates spread around the apical cell perimeter to form a tight junctional belt and seal the tissue (3). (B) Mesoscale events during tight junction belt formation. (1) ZO1 condensation leads to partitioning of junction proteins. (2) Adhesion of condensates to the apical interface via PATJ induces a wetting transition which drives spreading of isolated condensates along the apical interface into 1D belt. (3) Polymerization of tight junction strands in the ZO1 belt establishes the tight trans-epithelial barrier. (C) Molecular interactions underlying the wetting of ZO1 condensates along the apical interface. ZO1 condensates form at the lateral membrane around adhesion receptors (JAMA, CLAUDINs, NECTINs). PATJ binds to the apical membrane via the CRB complex (PALS1). Interactions of membrane bound ZO1 and PATJ drive adhesion of ZO1 condensates to the apical interface. Apical interface localization feeds back to ZO1 condensation and possibly actin and tight junction strand polymerization.

In addition to junctional condensation and wetting, belt formation involves active processes such as actin polymerization and claudin strand formation ^7,51^. Interestingly, our data suggests that there is feedback between adhesion of ZO1 condensates to the apical interface and these active processes. Deleting the apical adhesion domain in ΔL27-PATJ does not only prevent spreading of condensates along the apical interface but it also inhibits claudin polymerization and strand formation. Our live imaging data showed that in ΔL27-PATJ tissue nascent ZO1 condensates often dissolve after initial formation (Figure 2A-B), which suggests that interactions with the apical interface are required to maintain or even enhance the biophysical properties of ZO1 condensate. In addition, our FRAP measurements show that the remaining ZO1 membrane condensates in ΔL27-PATJ cells are less dynamic than in WT cells, which indicates a change in material properties (Figure 2C). Interestingly, ZO1 dynamics at the tight junction have been linked with actin cytoskeleton and ATP levels. In fact, blocking actin interactions by different drug treatments or depleting ATP levels significantly reduced ZO1 dynamics at tight junction^52,53^. Together, these observations suggest that wetting of ZO1 condensates to the apical interface feeds back to active processes at the interface, which can further enhance the wetting process.

A wetting-based mechanism of tight junction belt formation has several interesting implications. It explains the positioning and structure of the tight junction belt as a consequence of the collective physico-chemical properties of the condensed ZO1 scaffold. That apical polarity and the junctional complex are connected has been known for a long time^48^. However, how this connection relates to the assembly and the structure of the tight junction belt remained unclear. Wetting of junctional condensates provides a mechanism how assembly of cell adhesion complexes is guided by the cell polarity machinery and it directly bridges the molecular interaction scale to the mesoscale structure. Intriguingly, wetting of junctional condensates could also explain how the junctional complex can dynamically adapt to shape and length changes of the apical perimeter due to tissue mechanics. Any change in interface length due to stretching or contraction is directly “sensed” by the wetted ZO1 scaffold, which will spread thinner or thicker according to the interface length. How the collective properties of the wetted junctional condensates provide structurally flexibility and robustness to cell shape changes during development is an exciting avenue for future studies.

More generally, our work has important implications for how cells exploit the collective physical properties of protein condensates to actively shape higher order structures. Recent examples have shown that interactions of protein condensates with biological surfaces such as DNA/RNA, cytoskeletal filaments or lipid membranes can drive mesoscale shape changes such as DNA compaction, filament bundling and branching and membrane budding and wrapping ^8,10,16^. The common emerging theme in these different examples is that shape changes are driven by forces arising at the interface of protein condensates with cellular structures. Here we uncovered a striking example how cells use and modulate interfacial forces of membrane attached condensates to position and shape cell-cell adhesion complexes. We envision the emergent properties of protein condensates, can serve as general example how complex mesoscale structures self-assemble in cells. It also provides a new perspective on how to manipulate tight junction permeability in biomedical application.

## Supporting information

First hour of mN-ZO1 in WT tissue during junctional belt assembly

mN-ZO1 dynamics in WT cell-cell interface single junction belt assembly

mS-PATJ recruitment to mN-ZO1 cell-cell interface single junction belt assembly

## ACKNOWLEDMENTS

We thank the following MPI-CBG facilities: Genome engineering facility (Mihail Sarov and Ilka Reichardt) for generating the CRISPR-Cas9 L27-PATJ deletion, Cell Technologies (Julia Jarrells, Kathleen Kulb) for sequencing and qPCR, Protein expression (Aliona Bogdanova) for cloning. We thank Frank Stein (EMBL) for mass-spectrometry and help with bioinformatic analysis. We thank Andre Le Bivic for sharing the PATJ antibody. We thank Anthony A. Hyman for discussion and feedback. We thank Michele Marass for edits feedback.

This work was funded by the Max Planck Society and by the Deutsche Forschungsgemeinschaft (DFG) under project numbers 112927078 - TRR 83 and 402723784 - SPP2191. K.P.-G. was supported by an HFSP research grant (RGP0050/2018).

## AUTHOR CONTRIBUTION

A.H. and K.P-G conceived the project, wrote the paper and analysed the data. K.P-G performed all experiments except imaging of monolayers of Figure 2A and STED imaging done by A.H. and knock in cell lines done by C.M-L.

