## Supplementary Figures and Methods for "Wetting of junctional condensates along the apical interface promotes tight junction belt formation"

### Materials and Methods

#### Methods Cell Culture in 2D monolayers

MDCK-II (00062107, Public Health England) cells were cultured as a 2D monolayer in MEM with 5% FBS, 1% Non-essential amino acids (NeAA), 1% sodium pyruvate, 1% Glutamax without addition of antibiotics at 37 C with 5% $CO_2$. 2D monolayer on Transgenic cell lines were selected in presence of geneticin (400 ug/mL). Transient transfection of MDCK-II cells was carried out using Lipofectamine2000 after cells reach a confluency of 70%. Transgenic cell lines were created from the synthesized genes cloned via Not-I and Asc-I cutting sites into in house designed mammalian expression plasmids (pOCC series) with N-terminal Dendra-2, selection marker against Neomycin-Geneticin and a CMV promotor. To obtain stable lines, single clon selection was carried out via FACS.

Seeding of 2D monolayers on Transwell filters (Corning 3460), 400 µl of a 0.5 million cells per mL were transferred on the external side of a standing transwell filter. Cells were left to adhere at 37 ºC in a sterile environment for 30 minutes. Afterwards, each transwell filter was mounted into a 1- well plate with 1 mL of culture media on the bottom side and 500 µL of media on the internal side.

#### Generation of 3D Cyst and trans-epithelial permeability assay

For the adherent 3D cell culture cyst, MDCK-II cells were resuspended from a confluent monolayer into single cell suspension. The surface of the Mateck dish (355 mm glass bottom, P35G-0.170-14-C) was coated with a solution of laminin 0.5mg/ml for 1h at 37C 5%CO2. Afterwards, a suspension of 20k cells were seeded on the coated surface in the respective culture media complemented with in 5% Matrigel on ice. Cells were cultured forv5

-6 days until reaching 30 to 40 µm in diameter. To measure permeability of dextran (Alexa647, 10k) cyst medium was supplemented with 10µM dextran. After 15 min incubation cyst were imaged on a confocal microscope to evaluate distribution of dextran. Dextran permeability into the lumen was quantified by measuring the intensity in the lumen and dividing this to the average intensity outside the cysts.

**CRISPR/Cas9 knock-in MDCK-II cells**

The CRISPR-Cas9 technology was used to generate several single and double knock-in lines in MDCK-II cells. ZO1-mNeon was generated by fusing a copy of mNeon at the N-terminal of the initial exon of endogenous ZO1 targeting. By using that background cell line, several other scaffold proteins have been tagged with mScarlet at the N-terminal of the initial exon of endogenous PATJ, ZO2 and at the C-terminal for MAGI3 generating double knock in lines. Briefly, Specific CRISPR RNA (crRNA, IDT It-R® CRISPR-Cas9 crRNA) for each gen of interest were designed using online tools Crispor (http://crispor.tefor.net) and ChopChop (https://chopchop.cbu.uib.no). *Trans*-activating CRISPR RNA (tracrRNA, IDT catalog#1072532) and crRNA, were annealed at a ratio of 1:1 by incubating for 5 minutes at 95C and then 10 minutes at room temperature to generate gRNA. All gRNA can be found in key resource table. Next, the donor plasmid containing 5' and 3' homology arms were synthesised in a pUC57 Kan (Genescript) or a pUCIDT Kan (IDT). A PTisy plasmid containing the fluorescence tag (mSc or mNn) was integrated into the donor plasmid via digestion using restriction enzymes. Afterwards, RNP complex was assembled by mixing 1 ul (10 ug/ul) of the recombinant Alt-R® S.p. HiFi cas9 nuclease (IDT cataloq#1081060) and 1 ul of gRNA (100 uM) with the reaction buffer, followed by incubation at room temperature for 20 min. The RNP complex and 1 ug the donor plasmid was co-transfected *via* electroporation and overlap with the exon sides. Electroporation of each complex was performed in 300.000 cells (Invitrogen NEON electroporation machine and kit, 2 pulses, 20 ms, 1200 V). Cells are then plated into growth medium in a 6 well plate. Medium was exchanged after 24h. Cells were sorted 48-72h after electroporation. Fluorescent cells (mNn or mSc) were enriched (1 or 2 cycles) and single clones were plated into one well of a 96 well plate. To validate genetic modification, we first did a PCR amplification. To confirm the correct insertion of the fluorescence tag at the correct locus of the gen of interest was verified by sequencing genotyping.

**CRISPR/Cas9 knock-out in MDCK-II cells**

For the deletion of PATJ ∆Exon3, we used two guideRNAs on each site of the targeting exon for each gene were selected based on low off-target activity using http://crispor.tefor.net. The guideRNAs were ordered as crRNA from Integrated DNA Technologies (IDT). GuideRNAs were transfected in pairs and cell pools were tested for deletion events by PCR using primers spanning the targeting exon. Best performing gRNA pairs were used for cell cloning. Single cells were screened for the deletion by PCR using primers spanning the targeting exon. A flanking PCR with one primer outside and one primer inside of the deletion was performed on the knock-out candidate clones to verify the absence of the wild-type allele. Deletion alleles were verified by Sanger Sequencing.

**2-Colour Super-Resolution STED microscopy**

Confocal imaging was performed on a commercial confocal STED microscope (Abberior Instruments, Göttingen, Germany) with pulsed laser excitation (490 nm, 560 nm, 640 nm, 40MHz). 60x water and 100x oil objectives (Olympus). STED imaging was performed using a custom designed Abberior 775 3D-2 Color-STED system with 60×/1.2 NA Olympus water and 100×/1.4 NA oil Olympus objective. Star Orange was imaged with a pulsed laser at 560 nm, and excitation of Abberior Star Red was performed at 640 nm. The depletion laser for both colours was a 775 nm, 40 MHz pulsed laser (Katana HP, 3W, 1ns pulse duration, NKT Photonics). The optimal combination of excitation intensity, STED power and pixel dwell time were established by minimizing the onset of strong photo bleaching. To reduce high frequency noise, STED images were filtered with a 2D or 3D Gaussian with a sigma of 0.8 pixels. Quantification and segmentation of STED data was done using custom software written in Matlab.

**Live imaging microscopy**

Live-imaging kinetics was aquired in a Wide-Field Delta Vision Elite system equipped with an Olympus PlanApo N 60x / 1,42 objective, SSI-lumencor illumination system and Roper Evolve EMCCD camera. Perturbation experiments were acquired with a wide-field General Electric Delta Vision system equipped with an Olympus PlanApo N 60x / 1,42 objective, and mN & mCh was excited with Applied Precision Xenon Arc Lamp (V300-Y18).

**MDCK-II calcium switch assay**

Calcium depletion experiment was performed on a confluent monolayer of MDCK-II with the different cell lines. Cells were grown for 18 hours in calcium free media until the tight junctions were disrupted. For proteomics experiments, media containing calcium was added to the cells at 37C 5% $CO_2$ and each sample was prepared at 0, 0.5h, 1h, 3h,18h afterwards. For live imaging experiments, media containing calcium was added to the cells directly on the microscope chamber at 37C 5% CO2 and the tight junction formation was imaged every 1 minute over 3 hours or every 30 min for 18h.

#### Determination of tight junction length coverage per cell

To determine tight junction coverage length at tissue level (%) or per cell (□m/cell), we segmented by CellPose the cell periphery using the raw bright-field and segmented the TJ length using the ZO1-mNeon signal and skeletonize it (Fiji Plugin, skeleton) (and) normalize it to the cell number.
*%TJ coverage = (TJ length/Cell Periphery) x 100%*

#### Determination of tight junction protein recruitment kinetics

To determine ($t_{1/2}$) arrival kinetics of the mS-tagged tight junction proteins we segmented the condensed mN-ZO1 signal and quantified the mScarlet intensity in the segmented ZO1 condensates and the cytoplasm over time. Cell segmentation was done by using CellPose (https://github.com/mouseland/cellpose) and ZO1 condensates segmentation using the plugin Skeleton in FIJI. To directly correct for photo-bleaching artefacts, we calculate the ratio between the junctional and the cytoplasmic mScarlet signal for each time point. Assuming that bleaching is spatially homogenous the ratio is independent of bleaching and directly reports the enrichment of the protein in the condensed ZO1 (tight junction). Kinetic data was fitted using a HiIl slope model in Prism using a nonlin fit variable slope (four parameters) to calculate the EC50 of each protein.

#### Determination of ZO1 condensate length or eccentricity

Quantification of ZO1 condensate length was done on segmented ZO1 images using the function regionprops to measure the “MajorAxisLength” or “Eccentricity” of segmented condensates over time in MATLAB.

#### Fluorescence recovery after photobleaching (FRAP)

FRAP experiments in cells were carried out with the following settings. Region of interest (ROI) was bleached using a 405nm diode with 1.5 mW at the back focal plane of the objective with 100 ms pixel dwell time. Pre-bleach and post-bleach images were acquired with a 490 nm laser with 5 mW. Fluorescence recovery of mNeon was monitored for 1-20 min with a time resolution of 11 s. Cell movements during the recovery were corrected by registration of all frames to the first frame using the plugin StackReg in FIJI. To correct for photo-bleaching during the measurement. FRAP traces were evaluated and fitted in MATLAB.

**Immunoblotting and in-gel fluorescence**

Aliquots of ~ 5ug of input, elution and beads in lysate buffer (10% glycerol, 2% SDS, 1mM DTT, 1x Protease Inhibitor cocktail, 50 mM Tris-HCl, pH 8) were loaded onto a SDS PAGE 4- 20% Tris-Glycine Novex Gels were run at 90 mV for 3 hours. Transfer was done in an iBlot2 gel transfer system (ThermoFisher, IB21001) onto a nitrocellulose membrane (10 min, 20V). To detect protein levels by immunoblotting a iBind Flex system (ThermoFisher, SLF2000) was used and the following antibodies were used IRDye-800CW Streptavidin (1:4000, LI-COR, 92632230). As secondary antibodies used IRDye-800CW goat anti-mouse (1:4000, LI-COR, 92532210) and IRDye-680CW Goat anti-rabbit (925-68021, LI-COR). Membranes were scanned by LI-COR Odyssey system at 700 nm and 800 nm. In-Gel fluorescence of the tagged proteins (Dendra2, mNeon, mCherry) was done in a Thyphoon FLA 7000 scan system (GE).

**Immunofluorescence**

Fixation was performed in same way for both 2D monolayer and 3D cysts. Cells were fixed by 4% paraformaldehyde (PFA) in PBS for 10 min at room temperature, followed by quenching in 300mM glycine and permeabilized with 0.5% Trition X-100 in PBS for 10 min. Cells were blocked with 2% BSA and 0.1% Trition X-100 in PBS for 1h at rt. Staining of all primary and secondary antibodies were incubated at room temperature for 2 h or 30 min with a dilution of 1:100 or 1:200 respectably in blocking buffer. Staining for neutravidin-StarRed was performed at 1:1000 dilution for 1h at rt in blocking buffer.

**APEX2 labelling during tight junction formation**

Cells were plated on T75 flask after reaching confluency, full medium was substituted by medium without calcium for 18 hours for the cells to reach rounded non polarize state. Afterwards full medium was put back and APEX2 was performed at the different time points. APEX2 labeling was performed by adding 1mM Biotin phenol to the cells for 30 min before the addition for 1 minute of 1mM of $H_2O_2$. Immediately after, the reaction was quench in ice by washing 3 times with 1× PBS supplemented with 10 mM sodium ascorbate (Sigma A4034), 10 mM sodium azide, and 5 mM Trolox ((+/-)-6-Hydroxy-2,5,7,8-tetramethylchromane-2-carboxylic acid, Sigma 238813). To validate biotinylation via fluorescence microscopy, cells were immediately fixed after this step. For proteomics and western blot analyses, cells were lysed by scraping them off the growth surface in ice-cold lysis buffer (2% SDS, 10% Glycerol, 1mM DTT, 50 mM Tris-HCl, pH 8, 1xProtease Inhibitor Cocktail Set III EDTA-Free (EMD Millipore, cat no. 539134), supplemented with 10 mM sodium ascorbate, 10 mM sodium azide, and 5 mM Trolox. After collecting the lysate in low protein binding reaction tube, 5uL of benzonase were added and incubated in a shaker for 15 min at 37 C 700rpm. Finally, 1mM EDTA, 1mM EGTA were added and the sample was spin down in a table centrifuge to remove derbies. After lysis, protein concentration was checked with pierce 660-nm (Thermo Fisher Scientific, cat. no.22660) protein assay test before the pull down to ensure starting with the same protein concentration. Lysates were aliquot into 1500 □g of total protein snap frozen and store at -80C.

**Streptavidin pull-down of APEX2 biotinylated proteins**

All buffers used for pull-down proteomics experiments were freshly made and filter by 0.22um filter prior to use. Frozen lysates (1500 g protein) were diluted 1:10 in Tris 50mM pH8 reaching an incubation buffer of (0.2% SDS, 1% Glycerol, 1mM DTT, 1mM EGTA, 1mM EDTA in 50 mM TrisHCl, pH 8, 1x Protease Inhibitor). For affinity purification, ~ 100 L of streptavidin magnetic beads (Pierce, PI88817) were washed in incubation buffer two times before the binding. Beads were added to the sample in a total volume of 2mL and incubated for 2h at room temperature in a rotating wheel. Next, beads were pellet down by using a magnetic rack and the supernatant was collected and kept for further western blot analysis. Each sample of the beads containing the bind biotinylated proteins was washed with a series of ice-cold buffers (2mL each) to remove the unspecific binders. Twice with washing buffer (0.2% SDS, 1% Glycerol, 1mM DTT, 1mM EGTA, 1mM EDTA in 50 mM Tris-HCl pH 8, 1x Protease Inhibitor), 1 time with 1M KCl, 1 time with 2M Urea in 50 mM Tris-HCl pH 8, 1 time with 2 mM Biotin 50 mM Tris-HCl pH 8 and finally three times with 50 mM Tris-HCl pH 8. Biotinylated proteins were eluted from the beads by boiling the beads in 50uL of elution buffer (5% SDS, 10% Glycerol, 20mM DTT, in 50 mM Tris-HCl, pH 8, 1x Protease Inhibitor) at 95C for 10 min followed by cold down in ice and briefly spin down. Samples were placed on a magnetic rack and the eluate was collected to a new tube to send to proteomics and continue with SP3 method. Both magnetic beads and 10uL of eluate were together with the input loaded onto Western Blot for validation of the biotinylation experiments before MS.

**Mass Spectrometry and TMT labelling**

Reduction of cysteine containing proteins was performed with dithiothreitol (56°C, 30 min, 10 mM in 50 mM HEPES, pH 8.5). Reduced cysteines were alkylated with 2-chloroacetamide (room temperature, in the dark, 30 min, 20 mM in 50 mM HEPES, pH 8.5). Samples were prepared using the SP3 protocol [55] and 300 ng trypsin (sequencing grade, Promega) was added per sample for overnight digestion at 37°C. Peptides were labelled with TMT10plex [56] .Isobaric Label Reagent

(ThermoFisher) according the manufacturer's instructions. In short, 0.8 mg reagent was dissolved in 42 ul acetonitrile (100%) and 8 ul of stock was added and incubated for 1h rt. Quench with 5% hydroxylamine for 15 min. rt. Samples were combined for the TMT10plex and for further sample clean up an OASIS® HLB µElution Plate (Waters) was used. Offline high pH reverse phase fractionation done on an Agilent 1200 Infinity high-performance liquid chromatography system, equipped with a Gemini C18 column (3 µm, 110 Å, 100 x 1.0 mm, Phenomenex[57]. Offline high pH reverse phase fractionation was performed using an Agilent 1200 Infinity high-performance liquid chromatography (HPLC) system equipped with a quaternary pump, degasser, variable wavelength UV detector (254 nm), peltier-cooled autosampler, and fraction collector (both set at 10 °C for all samples).

The column was a Gemini C18 column (3 µm, 110 Å, 100 x 1.0 mm, Phenomenex) with a Gemini C18, 4 x 2.0 mm SecurityGuard (Phenomenex) cartridge as a guard column. The solvent system consisted of 20 mM ammonium formate (pH 10.0) (A) and 100% acetonitrile as mobile phase (B). The separation was accomplished at a mobile phase flow rate of 0.1 mL/min using the following linear gradient: 100% A for 2 min., from 100% A to 35% B in 59 min., to 85% B in a further 1 min., and held at 85% B for an additional 15 min., before returning to 100% A and re-equilibration for 13 min. Thirty-two fractions were collected along with the LC separation that were subsequently pooled into 6 fractions (depending on sample). Whereby the first and the two last fractions of the 32 were not used at all (discard). Pooled fractions were dried under vacuum centrifugation, reconstituted in 15 µL 1% formic acid, 4% acetonitrile for LC-MS/MS analysis.

**Mass Spectrometry analysis**

IsobarQuant and Mascot were used to process the acquired data, which was searched against the Uniprot Canis Lupus proteome database, which contains common contaminants and reversed sequences. The following modifications were included into the search parameters: Carbamidomethyl (C) and TMT10 (K) (fixed modifica- tions), Acetyl (N-term), Oxidation (M), and TMT10 (N-term) (variable mod-ifications). A mass error tolerance of 10 ppm and 0.02 Da was set for the full scan (MS1) and the MS/MS spectra, respectively. A maximum of two missed cleavages were allowed and the minimal peptide length was set to seven amino acids.

At least two unique peptides were required for protein identification. The FDR on peptide and protein level was set to 0.01. The R programming language (ISBN 3–900051–07–0) was used to analyze the raw output data of IsobarQuant. Potential batch effects were removed using the limma package. A variance stabilization normalization was applied to the raw data using the vsn package. Individual normalization coefficients were estimated for different time points compare to the non-calcium condition (t0). Normalized data were tested for differential expression using the limma package. The replicate factor was included into the linear model. For comparisons between different time points vs the t0 condition, hits were defined as those proteins with an FDR <5 % and a Fold change > 2. In different time point comparative experiments, proteins were first tested for their enrichment over −t0 control. The R package fdrtool35 was employed to calculate FDRs using the t values from the limma output. Proteins with an FDR < 5% and a consistent FC of at least 10% in each replicate were defined as hits. The ggplot2 R package was used to generate the graphical. False positive list based on non biotynolated proteins were removed.

**Quantification and statistical analysis**

Images were analyzed with FIJI (https://fiji.sc/) and MATLAB (Mathworks). All data are expressed as the mean ± the standard deviation (SD), mean ± the standard error of the mean (SEM), or mean ± 95% confidence intervals as stated in the figure legends and results. The value of n and what n represents (e.g., number of images, condensates or experimental replicates) is stated in figure legends and results. Two-tailed Student's t tests or one-ANOVA were used for normally distributed data, and Wilcoxon Rank-Sum tests were used for non-normally distributed data. A Pearson's Chi-square test was used to determine if data were distributed normally.

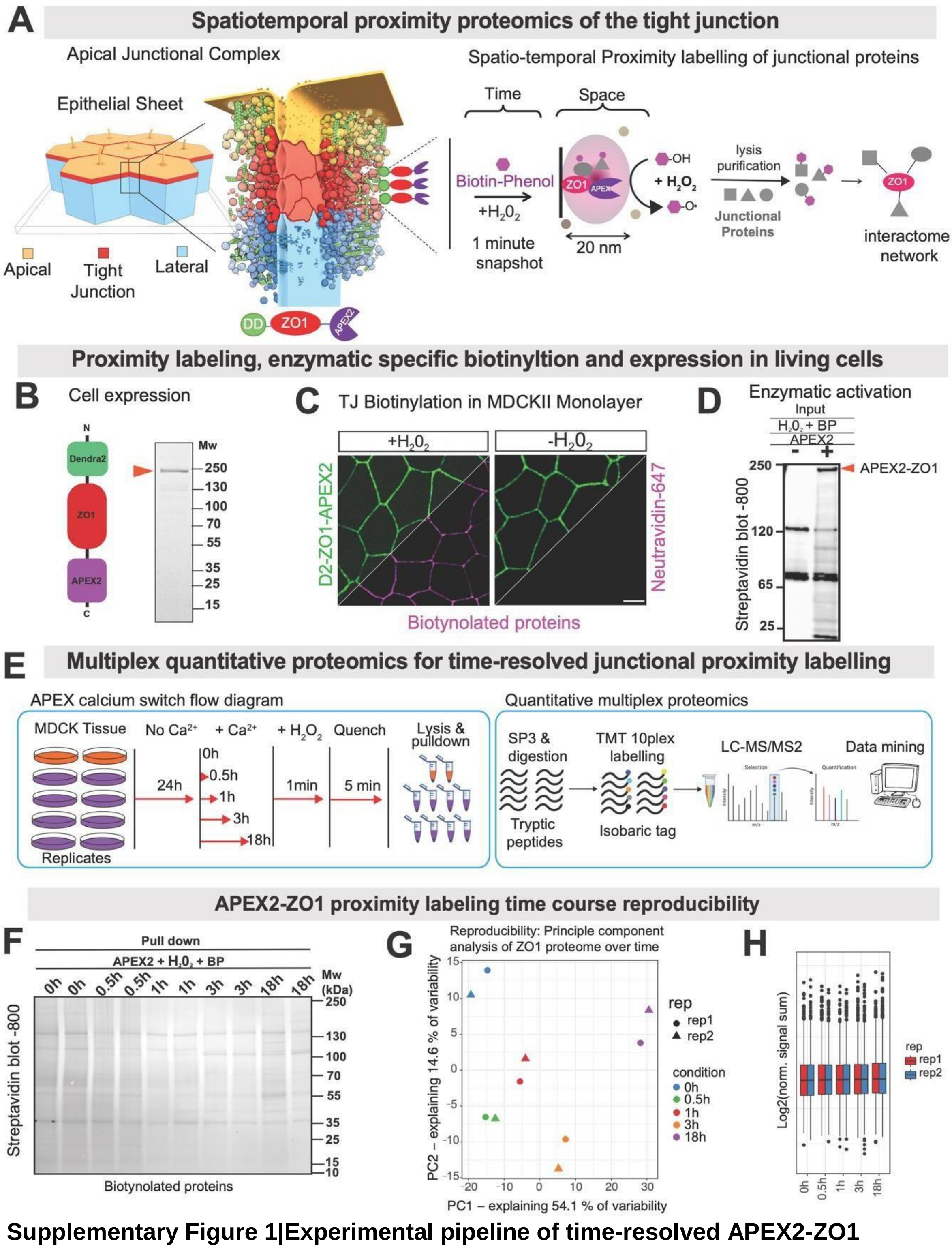


**Supplementary Figure 1|Experimental pipeline of time-resolved APEX2-ZO1 proximity proteomics.**

**Supplementary Figure 1|Experimental pipeline of time-resolved APEX2-ZO1 proximity proteomics.**

(A) Scheme of APEX2-ZO1 mediated protein labelling to identify protein interactions at the apical junctional complex. After activation for 1 min, proteins in proximity (< 20 nm) to the ZO1 scaffold get biotinylated and can subsequently be identified using mass spectrometry and develop protein networks.

(B) Dendra2-APEX2-ZO1 genetic construct and its expression on MDCK II cells with a MW ~ 250 kDa

(C) Correct localization of the D2-APEX2-ZO1 at the TJ-belt and confirmation of $H_2O_2$ specific activation and biotinylation of APEX2 labelling. Scale bar 10 µm.

(D) Activation of APEX2 and biotinylation pattern (+/- $H_2O_2$) analysed by western blot against streptavidin.

(E) Time-resolved APEX2-ZO1 proximity labelling was performed during the calcium switch assay at the four time points shown in Figure 1B. Biotinylated proteins were purified by streptavidin pull down, followed by SP3 enrichment, digestion and tandem mass tag labelling to allow quantitative mass spectrometry analysis.

(F) ZO1-APEX2 western blot image of the labelled biotinylated proteins after pull down at different time points used for proteomics

(G) Principal component analysis of ZO1 proximity proteome at different time points and replicates, showing that the proteome at different time points changed in a reproducible manner.

(H) Overall protein intensities across all samples at different time points and replicates show that protein content did not change.

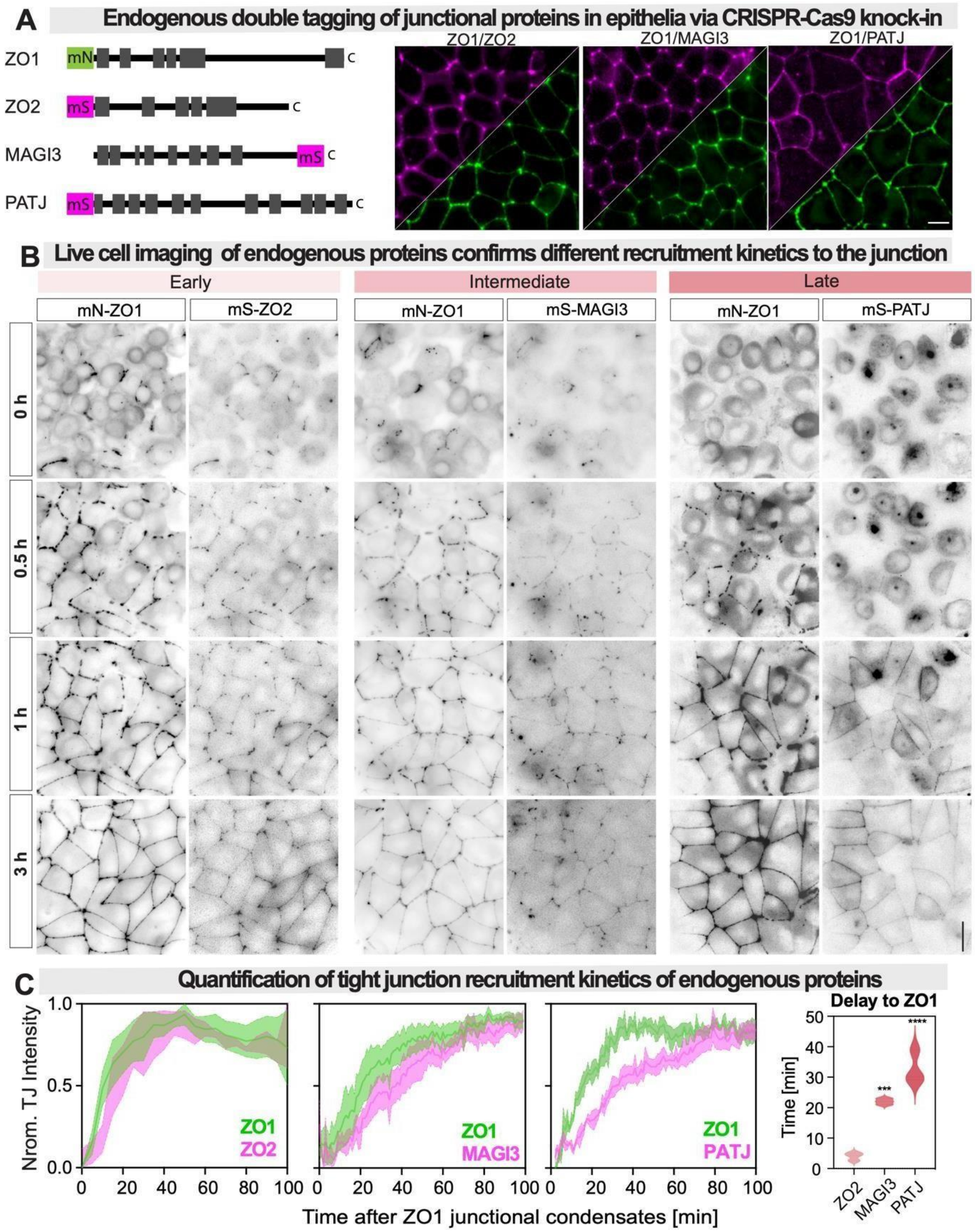


**Supplementary Figure 2|Live-imaging of endogenous proteins confirms recruitment kinetics of apical- junctional proteins to ZO1 condensates**

**Supplementary Figure 2|Live-imaging of endogenous proteins confirms recruitment kinetics of apical- junctional proteins to ZO1 condensates**

(A) Double fluorescence tagging via CRISPR/Cas9 mediated knock-in of mN-ZO1 (green) and mS-ZO2, mS-MAGI3 or mS-PATJ (pink) in MDCK-II cells. Double colour imaging of the three cell lines in confluent monolayers confirmed junctional co-localization of tagged proteins (see Figure S2A).

(B) Calcium switch imaging of the double tagged cell lines. Snapshots of same time steps as used for the APEX2 proximity proteomics are shown with inverted colour maps (0.5h,1h,3h). Scale bar 10 µm.

(C) Quantification of protein enrichment in mN-ZO1 condensates compared to the cytoplasm over time. For analysis pipeline see Figure S2B. Plots show the normalized enrichment of ZO2, MAGI3 and PATJ (magenta) at the tight junction compared to ZO1 (green). Kinetics were fitted with a Hill binding model to quantify the difference between the kinetics of ZO1 and other junctional proteins. The violin plot shows the difference in half-time of recruitment compared to ZO1. Significant comparison in reference to ZO2 (early) using a one-ANOVA test of n=3 experiments with cells > 100, ****P-value <0.0001.

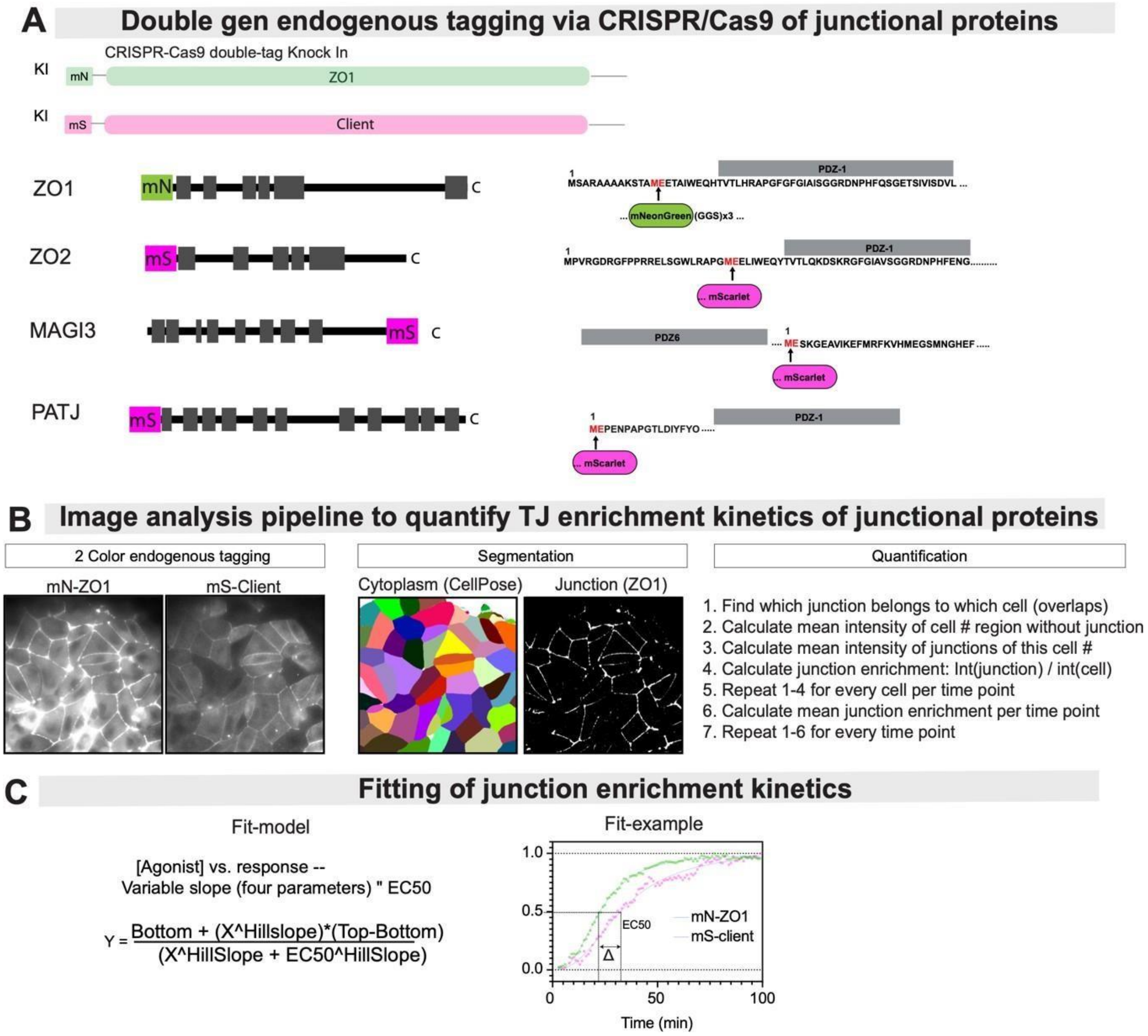


**Supplementary Figure 3|Quantification of recruitment kinetics of junctional proteins from live cell movies.**

(A) Endogenous tagging via CRISPR-Cas9 of three scaffold soluble junctional protein of each kinetic stage (ZO2, MAGI3, PATJ). Tagging with mNeon was used for ZO1 scaffold as a junctional reference and mScarlet for client protein.

(B) Image analysis pipeline showing time-resolved 2-color imaging of junction assembly. Cell cytoplasm was segmented using CellPose. Junction position was segmented using local intensity threshold of mN-ZO1. Junction enrichment was calculated locally for every cell as the ratio of junctional and cytoplasmic fluorescence intensity.

(C) Quantification of junction arrival time by fitting junction enrichment kinetics with Hill binding model. Delta between ZO1 and each client protein represents the different arrival times.

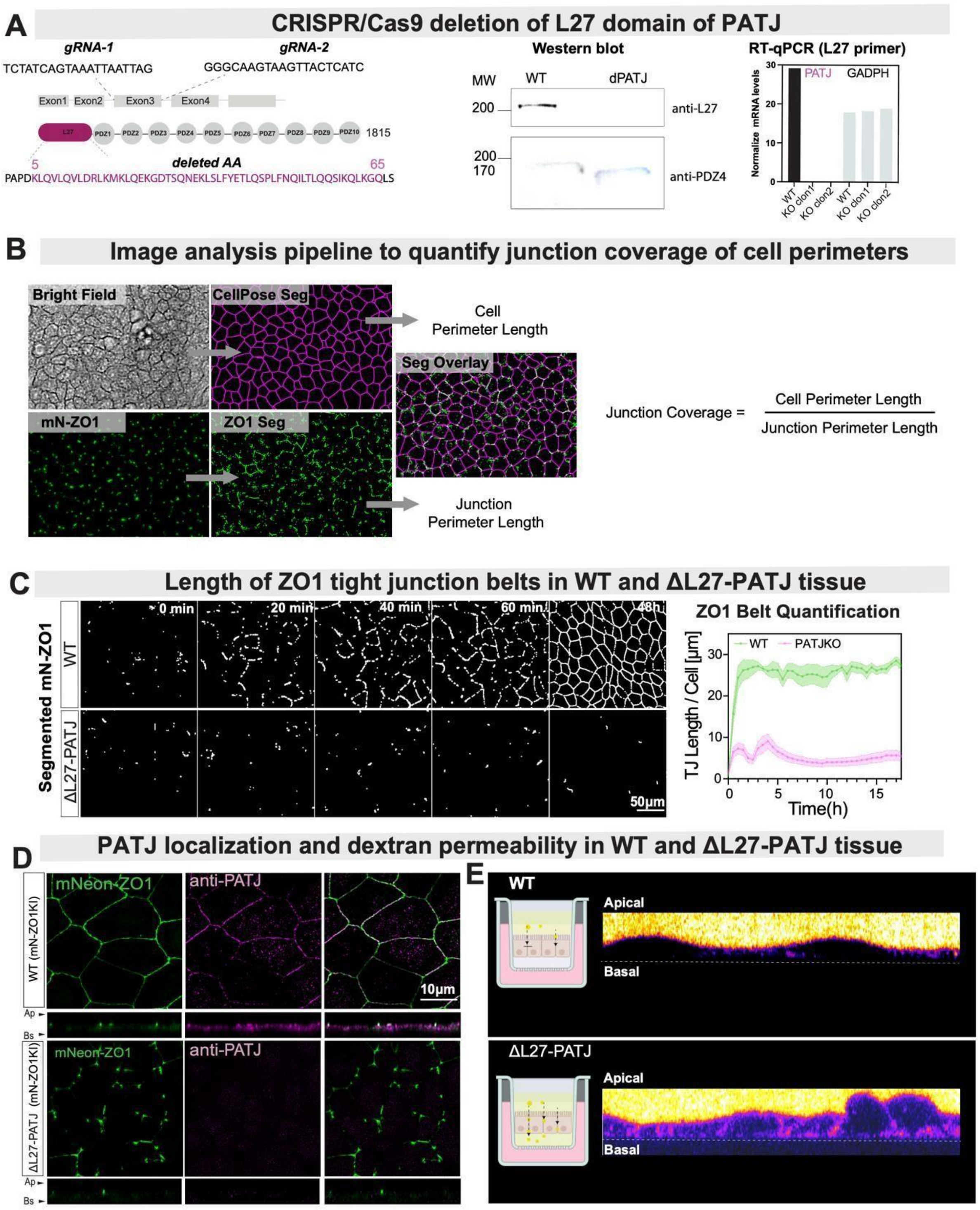


**Supplementary Figure 4|PATJ is required for tight junction formation and barrier function.**

**Supplementary Figure 4|PATJ is required for tight junction formation and barrier function.**

(A) Deletion of L27 domain of PATJ via CRISPR/Cas9 in background of mN-ZO1 knock in. Guides used to remove exon 3 and sequencing confirming the deletion. Middle panel Western Blot showing protein levels of PATJ WT vs KO for L27 domain and PDZ4 domain. Right panel qPCR showing the mRNA levels of PATJ WT vs two clones of the deleted L27 domain.

(B) Analysis pipeline of segmentation to quantify the % coverage of cell perimeters by mN-ZO1. Segmentation of the cell periphery was done using CellPose on phase contrast images. Segmentation of the ZO1-mN was done using local intensity thresholding and skeletonization in FIJI.

(C) Segmented skeletonize junctions over calcium switch assay to quantify TJ kinetics of WT vs ΔL27-PATJ MDCK cells over time (0-48h). Right panel shows the quantification of TJ length/cell after segmentation of WT and ΔL27-PATJ with completely different amplitude during 15h of TJ biogenesis. Graph represents mean +/-SEM over >100 cells in n=3.

(D) Staining of PATJ (pink) and ZO1(green) in WT vs ΔL27-PATJ in MDCK

(E) Trans-epithelial barrier assay measuring the diffusion of dextran molecules through MDCK monolayers in WT vs ΔL27-PATJ tissue seeded in transwell plates from apical side (yellow) to basal side (black)

**Supplementary Videos**

Video 1 – First hour of mN-ZO1 in WT tissue during junctional belt assembly

Video 2 – First hour of mN-ZO1 in ΔL27-PATJ tissue during junctional belt assembly

Video 3 – mN-ZO1 dynamics in WT cell-cell interface single junction belt assembly

Video 4 – mN-ZO1 dynamics in ΔL27-PATJ cell-cell interface single junction belt assembly

Video 5 – mS-PATJ recruitment to mN-ZO1 cell-cell interface single junction belt assembly

Video 6 – mS-PATJ recruitment to mN-ZO1 full tissue during junctional belt assembly
